# Fluorescein-based sensors to purify human α-cells for functional and transcriptomic analyses

**DOI:** 10.1101/2022.11.27.518097

**Authors:** Sevim Kahraman, Kimitaka Shibue, Dario F. De Jesus, Jiang Hu, Debasish Manna, Bridget K. Wagner, Amit Choudhary, Rohit N. Kulkarni

## Abstract

Pancreatic α-cells secrete glucagon, an insulin counter-regulatory peptide hormone critical for the maintenance of glucose homeostasis. Investigation of the function of human α-cells remains a challenge due to the lack of cost-effective purification methods to isolate high-quality α-cells from islets. Here, we use the reaction-based probe diacetylated Zinpyr1 (DA-ZP1) to introduce a novel and simple method for enriching live α-cells from dissociated human islet cells with > 97% purity. The α-cells, confirmed by sorting and immunostaining for glucagon, were cultured up to 10 days to form α-pseudoislets. The α-pseudoislets could be maintained in culture without significant loss of viability, and responded to glucose challenge by secreting appropriate levels of glucagon. RNA-sequencing analyses (RNA-seq) revealed that expression levels of key α-cell identity genes were sustained in culture while some of the genes such as *DLK1, GSN, SMIM24* were altered in α-pseudoislets in a time-dependent manner. In conclusion, we report a method to sort human primary α-cells with high purity that can be used for downstream analyses such as functional and transcriptional studies.

## Introduction

Zinc binding molecules such as Newport Green (1) and ZIGIR (zinc granule indicator) (2) have been employed previously for the purification of human or murine islet cells. We have recently reported the purification of human pancreatic β-cells (3) and stem-cell derived β-like cells (4) using the zinc-based reaction probe DA-ZP1. DA-ZP1 is a non-fluorescent zinc sensor that binds Zn(II) with nanomolar affinity (5). Binding of Zn(II) selectively and rapidly mediates hydrolytic cleavage of the acetyl groups and generates a strong fluorescence signal that can be used to sort the labelled cells by fluorescence activated cell sorting (FACS). In Lee *et. al*., we performed FACS analysis of DA-ZP1 stained human islets cells and observed enrichment of β-cells in the DA-ZP1 positive population and labeled the remainder of the cells DA-ZP1 negative (3). In the present study, we investigated whether zinc probes can also be used to identify a purified population of α-cells in a mixed population of pancreatic endocrine cells. We classified DA-ZP1 positive cells as either “DA-ZP1 intermediate” or “DA-ZP1 bright” by distinctly marking the boundary among the fractions to contrast those with “bright” DA-ZP1 fluorescence that were identified as β-cells. These data indicate that high purity α-cells and β-cells can be generated by simultaneously sorting islet cells using a zinc reaction probe (DA-ZP1). This approach could be useful for studying purified primary human α-cells for functional analyses and for comprehensive analysis of the transcriptomes to further increase our understanding of α-cell biology in health and disease.

## Results

### DA-ZP1 as a tool to purify live human pancreatic α-cells

To test whether DA-ZP1 is able to sort pancreatic α-cells from a mixed population of pancreatic endocrine cells, we dispersed human islets into single cells and labeled them with DA-ZP1 (Fig. 1a). Flow cytometry analysis of the cells showed a wide spread of fluorescence intensity among dispersed islet cells on a twodimensional density plot. We classified the cells into three subsets based on their fluorescence intensity and drew a gate to separate each of the subsets (Fig. 1b). The subset centered near the unstained cell background showed ‘low’ fluorescence intensity while the other two cell populations positioned on the right side of the dot-plot showed ‘intermediate’ and ‘bright’ fluorescence intensities, respectively (Fig. 1a, b). We confirmed that DA-ZP1 labeling resulted in a similar fluorescence intensity pattern among the three cell populations (low, intermediate, bright) in an additional three independent human islet donors (Supplementary Fig. 1a, b). To identify percentage of hormone-containing cells in each subset, the sorted cells were plated, fixed, and immunostained using antibodies to detect insulin or glucagon. Immunofluorescence analysis showed that the bright subset was highly enriched for human beta cells (~ 83% CPEP+ cells), while the intermediate subset consisted of α-cells of high purity (~ 97% GCG+ cells) (Fig. 1c, d). The cells in the DA-ZP1 ‘low’ subset were mostly hormone negative, indicating that these cells likely represented non-hormonal cells such as fibroblast-like, endothelial, or exocrine cells. Consistent with the FACS-based fluorescence assessment, fluorescent microscopy validated that the DA-ZP1 ‘bright’ subset displayed higher fluorescence signal compared to the intermediate subset, and that the unsorted islet cells are comprised of a mixture of cells with varying fluorescence intensities (Fig. 1e, Supplementary Fig. 1c).

**Figure 1.**
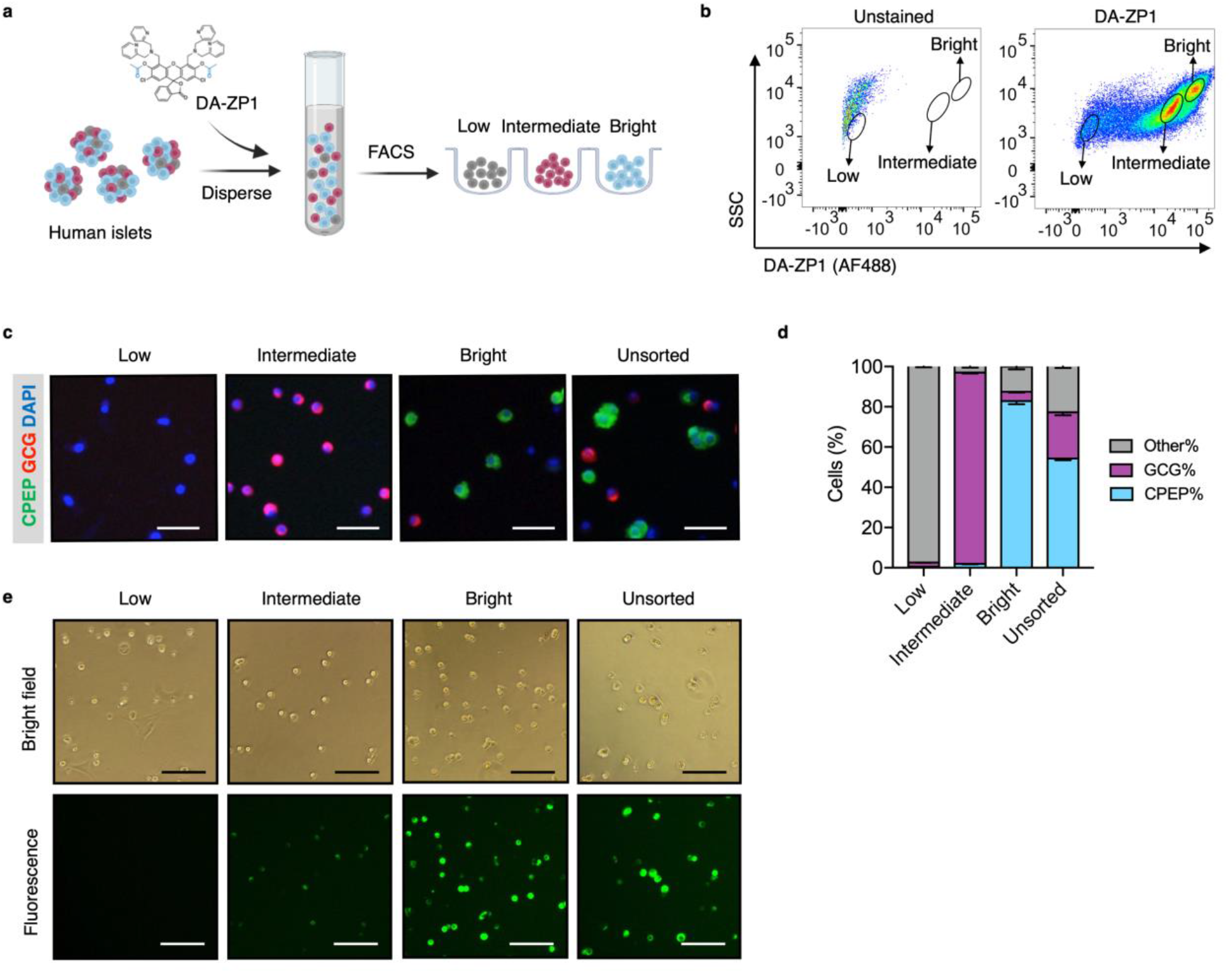
Isolation of live human pancreatic α-cells after staining with DA-ZP1 by FAC-sorted method. **a,** Experimental outline. **b,** Representative FACS plot showing three cell populations with low, intermediate, or bright fluorescence. The plot represents the data collected from Donor-1 islets. Unstained (left) vs DA-ZP1 treated (right) human islets. Gating strategy and the data collected from the other donors (n = 3) are given in Supplementary Fig. 1. **c,** Representative images of human islet cells after FAC-sorting showing C-peptide (green) and glucagon (red) expressing islet cells. Nuclei were stained with DAPI (blue). Scale bar, 50 m. **d,** Quantification of percentage of CPEP+, GCG+, and other cells (CPEP-GCG-) in each cell population. Data are presented as mean values ± s.e.m. **e,** The DA-ZP1 derived green fluorescence is maintained in the next day of sorting in the sorted islet cells. The cells were plated in Matrigel-coated flat-bottom plates. Scale bar, 100 μm. n = 3 biological replicates using islet cells from a single donor. **(b-e)**. See also Supplementary Fig. 1c.

### α-Pseudoislets can be maintained in culture without losing their viability

Purified pancreatic α-cells were maintained in culture for the assessment of viability post-sorting (Fig. 2a). Both the sorted α-cells and the unsorted islet cells formed islet-like clusters shortly after seeding in U-bottom low attachment plates (Fig. 2b, c). We refer to DA-ZP1 intermediate cells as α-pseudoislets, since they consisted of highly purified GCG+ cells and formed islet-like cell clusters. The size of the clusters was proportional to the number of cells in each well. Interestingly, α-pseudoislets tended to form tighter clusters shortly after plating compared to unsorted islet cells (Supplementary Fig. 2). Cell viability was determined by measuring intracellular ATP levels on day 5 and day 10 post-sorting. Culturing cells for up to 10 days did not alter viability of α-pseudoislets and unsorted islet cells indicating that these cells can be maintained in culture without significant cell loss. In contrast, intact whole islet cells started to die after day 5 possibly due to necrosis in the core of the islets caused by hypoxia (13,14) (Fig. 2d). We further validated that α-pseudoislets consisted of a highly pure population of α-cells with > 97% GCG+ cells on day 5 and 10 (Fig. 2e, f). To assess α-cell death, apoptotic index was measured by quantification of the percentage of TUNEL+GCG+ cells. We detected a small number of apoptotic GCG+ cells on day 5 (0.05% ± 0.02) and on day 10 (0.09% ± 0.04) in α-pseudoislets. In contrast, the apoptotic index of α-cells in unsorted islet cells was higher than that of α-pseudoislets on days 5 and 10 (Fig. 2g, h). Apoptosis index remained stable over the duration of the culture in both α-pseudoislets and in unsorted islet cells indicating that α-cells survived even after they form pseudoislets. However, the percentage of apoptotic α-cells in whole islets increased significantly on day 10 compared to day 5 which was likely due to hypoxia (13). Notably, we observed proliferating α-cells in α-pseudoislets, unsorted islet cells, and whole islets on both days 5 and 10 (Fig. 2i). Percentage of Ki67+GCG+ cells was similar in each group on day 5 (α-pseudoislets; 0.038% ± 0.008, unsorted islet cells; 0.049% ± 0.007, whole islets; 0.031% ± 0.004) and did not alter significantly with time spent in culture indicating that α-cells maintained their proliferation potential in culture (Fig. 2j). The relevance of this interesting observation requires further investigation.

**Figure 2.**
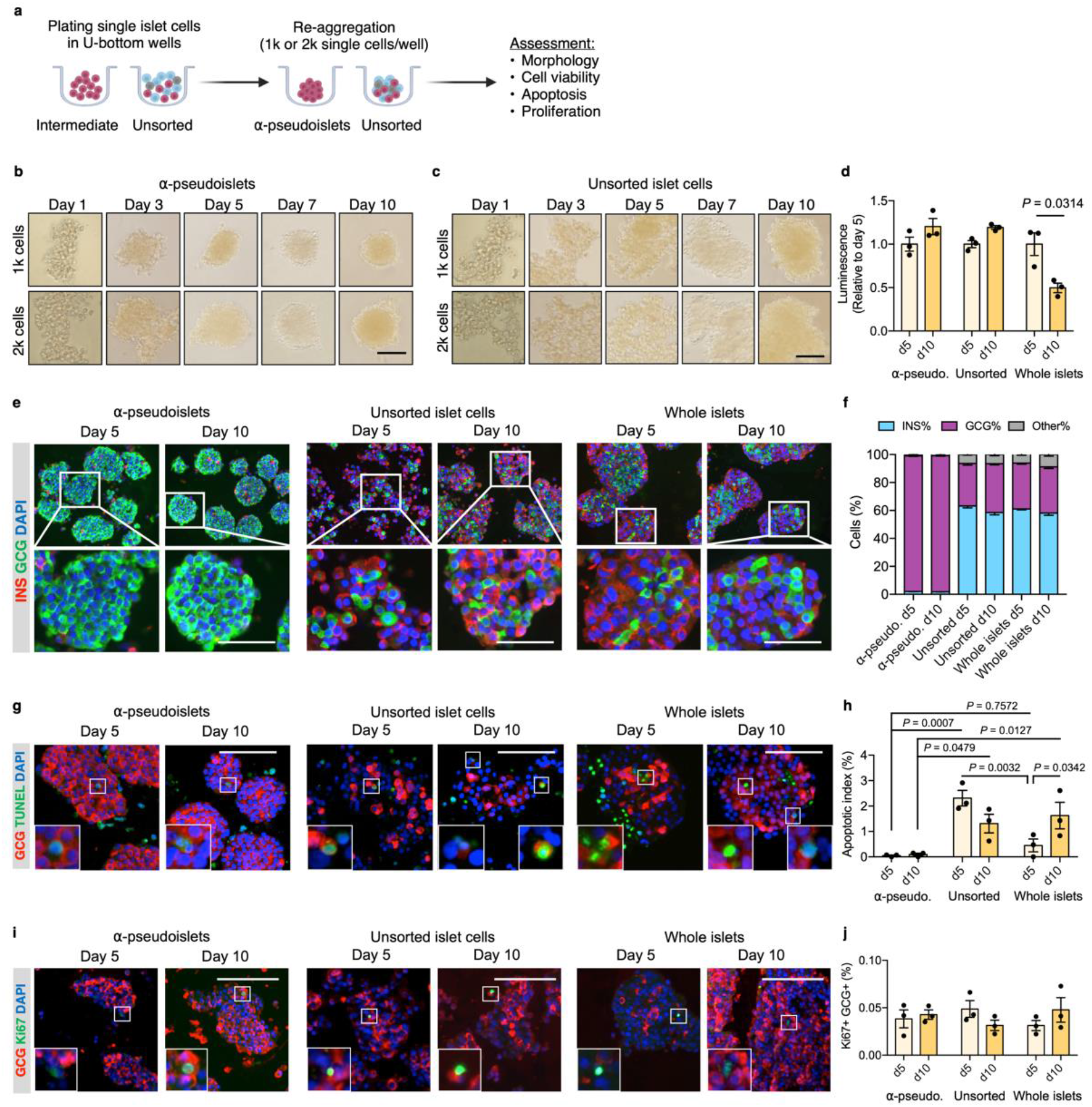
α-Pseudoislets are viable and able to function in vitro post-sorting. **a,** The single islet cells were seeded in U-bottom wells (1k or 2k cells per well) after sorting to allow re-aggregation. **b,c,** Bright field images of sorted α-cells **(b)** and unsorted islet cells **(c)** post-sorting. 1k (top panel) or 2k (bottom panel) single cells were seeded per well. Scale bar, 100 μm. See also Supplementary Fig. 2. **d,** Cell viability was quantified by luminescence reflecting intracellular ATP levels on days 5 and 10 following FAC-sorting. Fold-change relative to day 5. **e,** Representative immunostaining images of α-pseudoislets, unsorted islet cells, and whole islets on day 5 and day 10 showing INS (red), GCG (green). Nuclei stained with DAPI are blue. Scale bar, 100 μm. **f,** Percentage of INS+, GCG+, and other (INS-GCG-) islet cells. **g,** Representative immunostaining images of α-pseudoislets, unsorted islet cells, and whole islets on day 5 and day 10 showing GCG (red), TUNEL (green). Nuclei stained with DAPI are blue. Scale bar, 100 μm. Boxes show apoptotic α-cells. **h,** Percentage of TUNEL+GCG+ cells. **i,** Representative immunostaining images of α-pseudoislets, unsorted islet cells, and whole islets on day 5 and day 10 showing GCG (red), Ki67 (green). Nuclei stained with DAPI are blue. Scale bar, 50 μm. Boxes show proliferating α-cells. **j,** Percentage of Ki67+GCG+ cells. Data are presented as mean values ± s.e.m **(b-j)**. n = 3 biological replicates using islet cells from a single donor **(b-j)**. Two-way ANOVA followed by Sidak’s multiple comparison test **(d,h,j)**.

### α-Pseudoislets showed glucose-responsive glucagon release

Next, to investigate functional integrity, α-pseudoislets, unsorted islet cells, and whole islets were each independently challenged with either low (3.3 mM) or high (16.7 mM) glucose after preincubation in 16.7 mM glucose (Fig. 3a). Glucagon release significantly decreased in response to high glucose treatment in each group (Fig. 3b), while the fold increase between low (3.3 mM) or high (16.7 mM) glucose were comparable among the groups (Fig. 3c). Unsorted islet cells displayed higher basal secretion of glucagon compared to whole islets which is consistent with the previous observation (15). α-Pseudoislets showed higher glucagon content compared to unsorted islet cells (Fig. 3d), indicating high purity of sorted glucagon positive cells. These results indicate that α-pseudoislets are similar to unsorted islet cells in their capacity for glucose-responsive glucagon secretion.

**Fig. 3.**
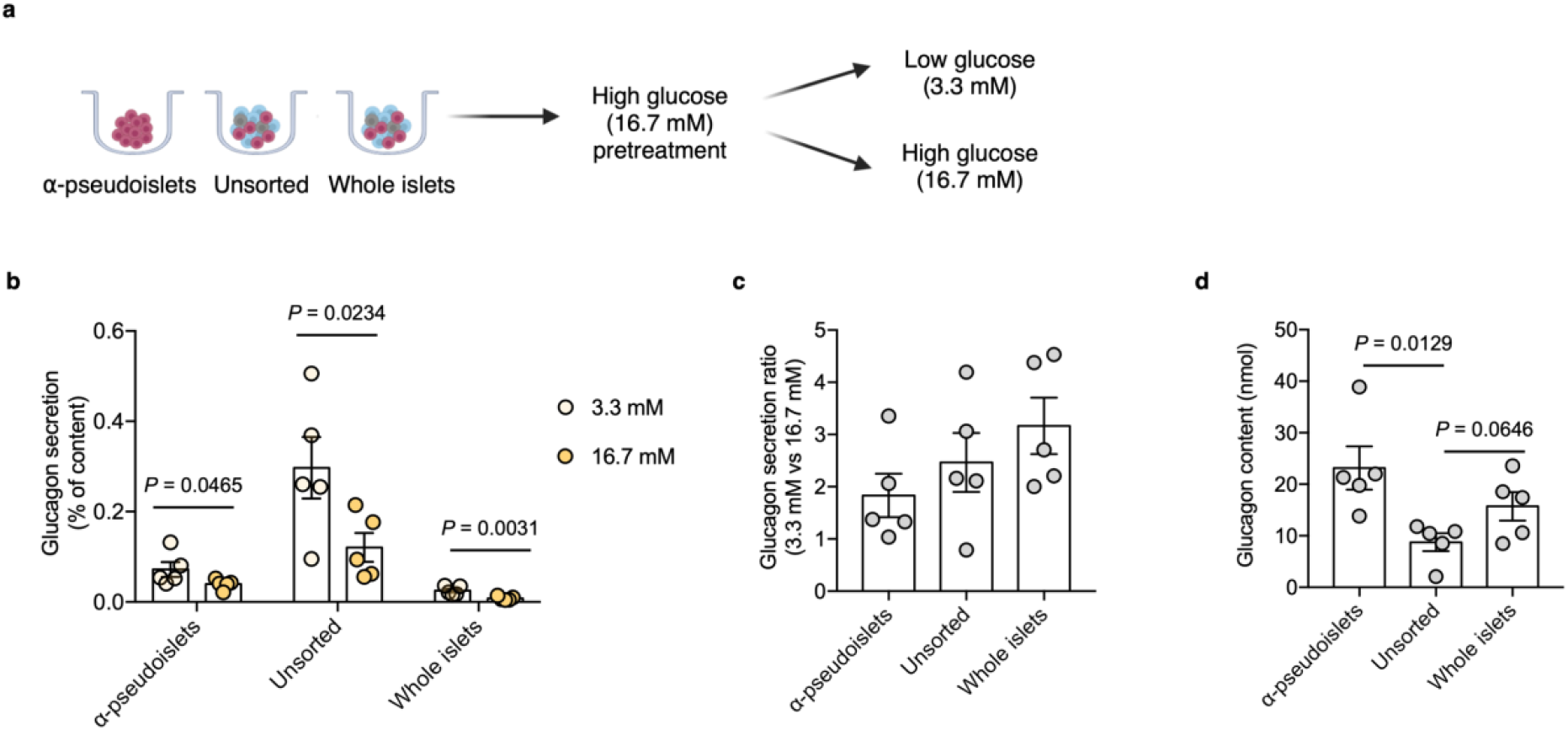
Glucagon secretion in response to glucose challenge. **a,** α-Pseudoislets, unsorted islet cells, and whole islets were pre-incubated in KRB buffer with 16.7 mM glucose followed by the incubation in KRB buffer with 3.3 mM glucose and 16.7 mM glucose. **b,** Glucagon secretion in response to glucose challenge (3.3 mM vs 16.7 mM). One-tailed Student’s t-test. **c,** Ratio of glucagon released by each groups of cells at 16.7 mM glucose versus that at 3.3 mM glucose. **d,** Glucagon content measured in each well containing ~8,000 cells (8 α-pseudoislets, 8 unsorted islet cells, and 8 IEQ). Data are presented as mean values ± s.e.m. **(b-d)**. n = 5 biological replicates using islet cells from a single donor **(b-d)**. Two-tailed multiple t-test corrected using Holm-Sidak method applied to **c,d**.

### α-Pseudoislets maintain α-cell identity in culture

To explore whether α-pseudoislets maintain α-cell identity by preserving genes enriched in α-cells, we compared the RNA-seq data from α-pseudoislets, unsorted islet cells, and whole islets on days 0, 5, 10. We first used a publicly available single cell RNA-seq (scRNA-seq) database performed on cadaveric human islets (GSE84133) (11) to identify genes that are differentially expressed between cadaveric islet α-cells and β-cells (α-cell enriched and β-cell enriched genes). Analysis of scRNA-seq data revealed 75 α-cell enriched genes and 68 β-cell enriched genes in cadaveric islet α-cells and β-cells, respectively [false discovery rate (FDR) < 0.1, fold change (FC) > 1.5, Supplementary File 1]. Expression analysis of α-cell enriched and β-cell enriched genes in α-pseudoislets showed that forty-two (56%) of the 75 α-cell enriched genes were upregulated and 27 (40%) of the 68 β-cell enriched genes were downregulated in α-pseudoislets compared to unsorted islets cells or whole islets on day 0 (FDR < 0.1; FC > 2 or FC < −2, Supplementary File 2), which confirms the enrichment of α-cells in α-pseudoislets (Fig. 4a, Supplementary Fig. 3). Concurrently, fifty-two (~ 70%) of the 75 α-cell enriched genes did not alter in α-pseudoislets on day 5 and day 10 compared to day 0 (FDR < 0.1; FC < −2 and FC > 2, Supplementary File 3, Fig. 4b), indicating the majority of the α-cell enriched genes including *GCG, MAFB, ARX, IRX2*, and *TTR* were preserved in culture. Similarly, fifty-three (71%) of the 75 α-cell enriched genes and 49 (72%) of the 68 β-cell enriched genes did not alter in unsorted islet cells on day 5 compared to day 0. Expression levels of the altered genes returned to normal levels on day 10 which indicated that existence of other islet cells and paracrine interactions were important for maintenance of α-cell or β-cell identity (16) (Fig. 4b, Supplementary File 3). On the other hand, whole islets did not show any changes in expression levels of α-cell enriched or β-cell enriched genes on days 5 and 10 compared to day 0 indicated that maintenance of islet structure was likely necessary to maintain cell identity. In sum, comparison of α-pseudoislets, unsorted islet cells, and whole islets on days 0, 5, 10 showed that co-existence of other cells and intact islet architecture were desirable but not indispensable for maintenance of α-cell identity *in vitro*.

**Fig. 4.**
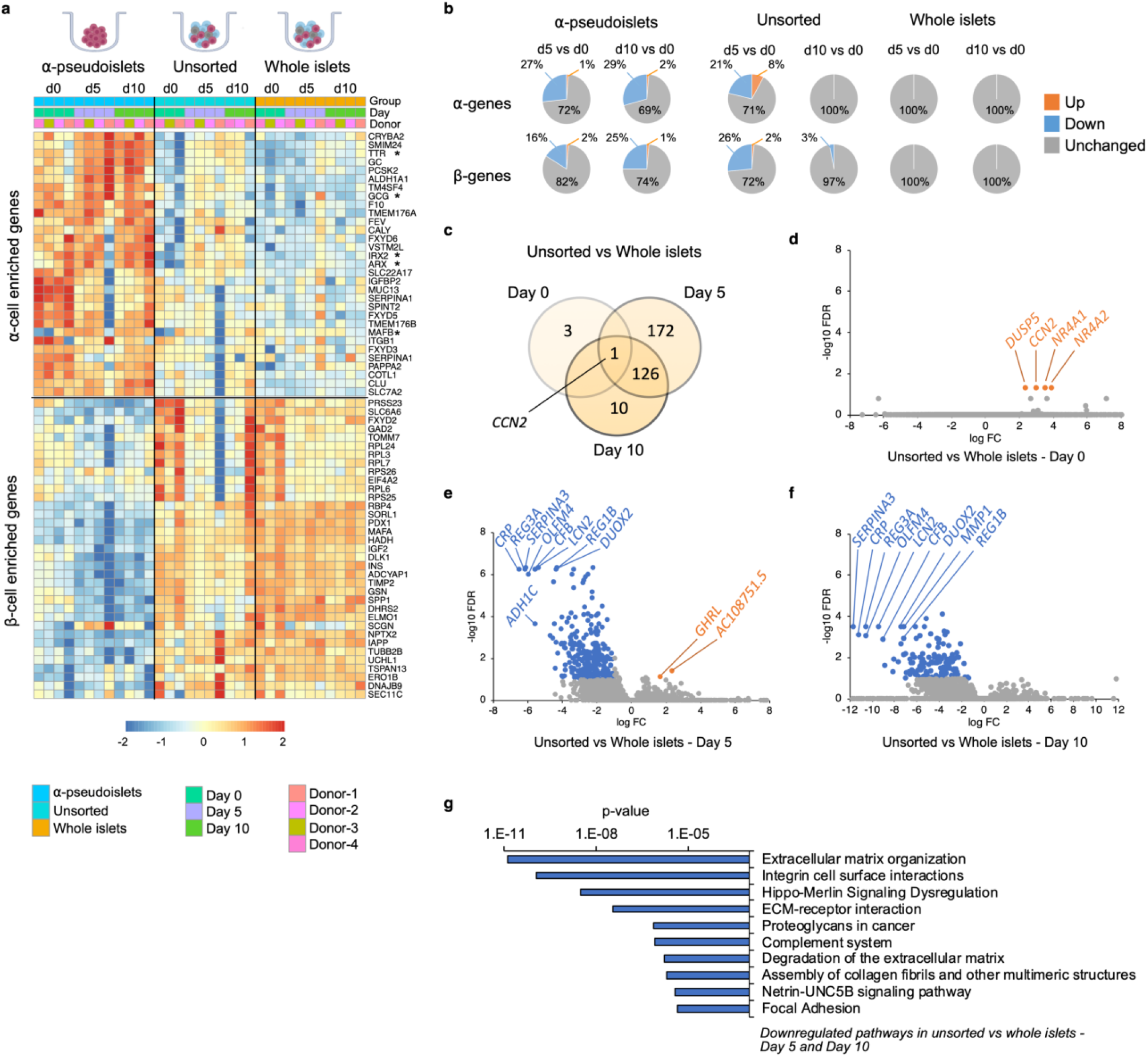
Changes in gene expression levels driven by dissociation and re-aggregation of human islet cells. **a,** Heatmap showing expression levels of α-cell enriched and β-cell enriched genes in α-pseudoislets, unsorted islet cells, and whole islets on day 0, 5, and 10. Asterisks show genes associated with α-cell identity and function (GCG, MAFB, ARX, IRX2, TTR). **b,** Pie charts showing percentage of α-cell enriched (top panel) and β-cell enriched (bottom panel) genes that alter in α-pseudoislets, unsorted islet cells, and whole islets on day 5 or day 10 compared to day 0. See also Supplementary Fig. 3. **c,** Transcriptome of unsorted islet cells was compared with whole islets on days 0, 5, 10. Venn diagram shows number of DEGs between unsorted islets cells and whole islets on different days. **d-f,** Volcano plots showing genes downregulated (blue) or upregulated (orange) significantly (FC < −2 or FC > 2, respectively, FDR < 0.1) on day 0 **(d)**, day 5 **(e),** and day 10 **(f)**. Gray shows non-significant genes with FDR > 0.1 and −2 < FC < 2. **g,** Top 10 pathways down regulated in unsorted islet cells on day 5 and day 10 compared to whole islets. n = 4; α-pseudoislets d0, d5, d10, unsorted islet cells d5, whole islets d5, d10, and n = 3; unsorted islet cells d0, d10, whole islets d0 using islets from four donors **(a-g)**.

### Extracellular matrix (ECM) organization genes are downregulated in dissociated and re-aggregated islet cells

To investigate the potential transcriptional changes that occur secondary to cell-to-cell interactions, we compared the transcriptome of whole islets with that of unsorted islet cells on days 0, 5, and 10 of culture (Fig. 4c). We filtered differentially expressed genes (DEGs) between unsorted islets cells and whole islets on days 0, 5, and 10 (FDR < 0.1; FC > 2 or FC < −2; Supplementary File 4). Differential expression analysis yielded 4, 299, and 137 DEGs on days 0, 5, and 10, respectively (Fig. 4c). Not surprisingly, the gene expression profile of unsorted islet cells was almost identical to whole islets on day 0 (Fig. 4d). For example, we detected differences in expression levels between the groups for only 4 genes, namely, *CCN2* (Cellular Communication Network Factor 2), *DUSP5* (Dual Specificity Phosphatase 5), *NR4A1* (Nuclear Receptor Subfamily 4 Group A Member 1), *NR4A2* (Nuclear Receptor Subfamily 4 Group A Member 2) on day 0 (Fig. 4d). All 4 genes were significantly upregulated in unsorted islet cells which indicates that dissociation of whole islets into single cells triggered acute changes in their expression. We detected changes in expression levels of 299 genes on day 5, among which 297 were downregulated and 2 (*GHRL, AC10875.5*) were upregulated in unsorted islet cells compared to whole islet cells (Fig. 4e). On day 10, all the 137 genes that were altered significantly in unsorted islet cells compared to whole islets were downregulated (Fig. 4f). Among these, 127 genes were consistently downregulated on day 5 as well as day 10. Pathway analysis of commonly downregulated genes on days 5 and 10 in the unsorted islet cells compared to whole islets showed changes in pathways such as ECM organization, integrin cell surface interactions, degradation of the ECM, complement system, and focal adhesion supporting the notion that the differences were likely due to physical separation of islets into single cells (Fig. 4g, Supplementary File 5). Interestingly, we observed consistent downregulation of genes involved in the Hippo signaling pathway such as *TEAD2, YAP1, WWTR1, TGFB2, CCN2* on days 5 and 10. Expression of *CCN2* showed a dynamic change with upregulation on day 0 and downregulation on days 5 and 10 in unsorted islet cells compared to whole islets.

### Time-dependent changes in the α-pseudoislet transcriptome

Next, to investigate how the gene expression pattern of α-cells alters after separation from neighboring non-α islet cells, we explored genes and pathways that are progressively up or downregulated in α-pseudoislets, unsorted islet cells, and whole islets in culture over the period from day 0 to day 10. We identified 413 genes in α-pseudoislets, and 341 genes in unsorted islet cells, in contrast to only 11 genes in whole islets that were significantly altered during this period (Fig. 5a, Supplementary File 6). The fact that the majority of the genes (597 out of 608 genes) that were altered during this period in α-pseudoislets and unsorted islet cells but not in whole islets likely reflects a transcriptional response of the cells following cell dissociation and reaggregation. Pathway analyses of these 597 genes that were altered in α-pseudoislets or in unsorted islet cells but not in whole islets, revealed networks such as ECM organization, integrin cell surface interaction, focal adhesion, and collagen formation (Fig. 5b, Supplementary File 7).

**Fig. 5.**
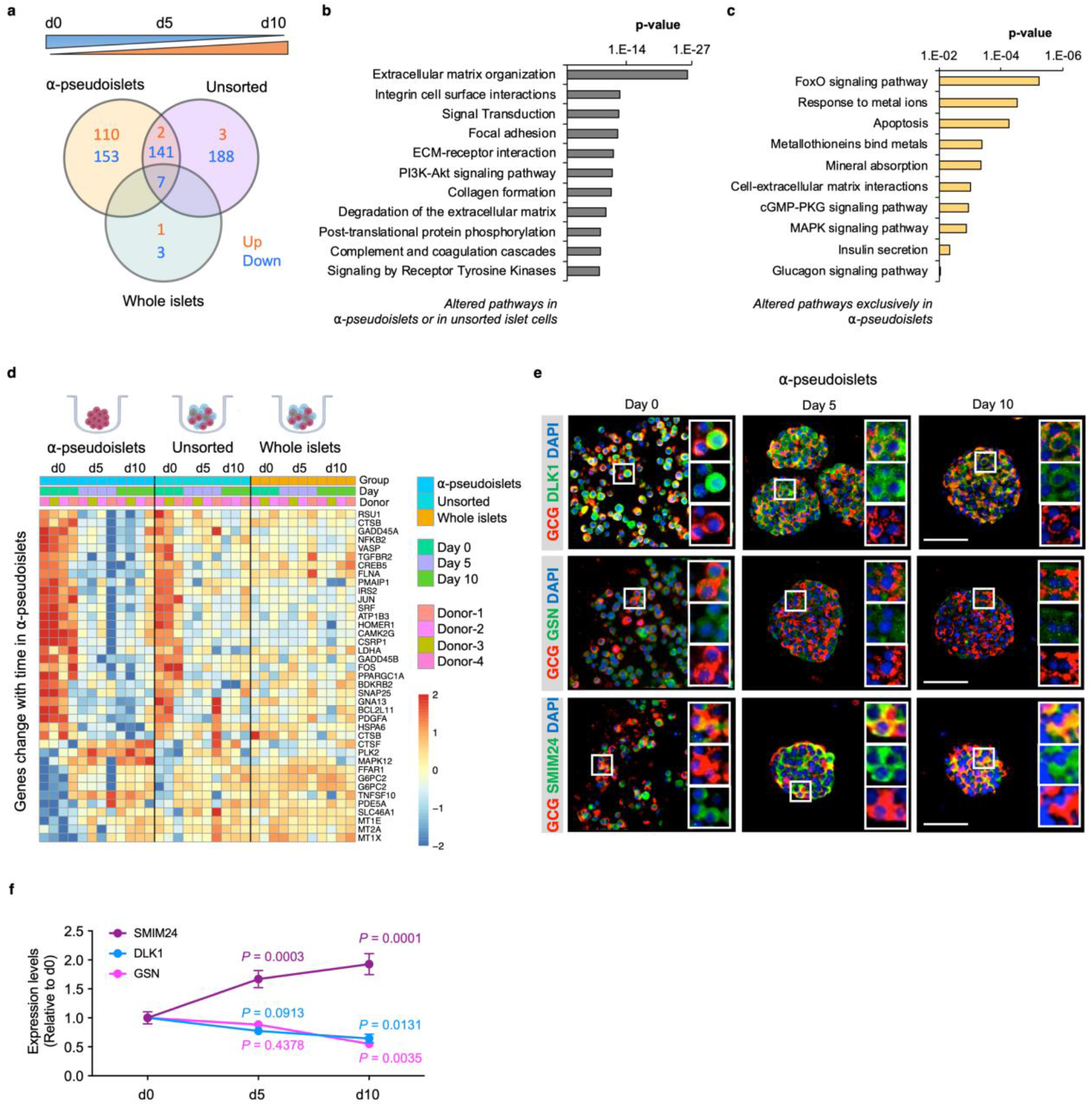
Time-dependent changes in transcriptome of α-pseudoislets. **a,** Venn diagram shows number of genes that progressively up or down regulated in α-pseudoislets, unsorted islet cells, and whole islets in culture over the period from day 0 to day 10. **b,c,** Pathway analysis showing altered pathways in α-pseudoislets or in unsorted islet cells except whole islets **(b)**, and altered pathways only in α-pseudoislets with time **(c)**. **d,** Heatmap showing expression levels of timeresponse genes altered only in α-pseudoislets. n = 4; α-pseudoislets d0, d5, d10, unsorted islet cells d5, whole islets d5, d10, and n = 3; unsorted islet cells d0, d10, whole islets d0 using islets from four donors **(a-d)**. **e,** Representative immunostaining images of α-pseudoislets on day 0, 5, and 10 showing GCG (red), DLK1, GSN, and SMIM24 (green). Nuclei stained with DAPI are blue. Scale bar, 50 μm. **f,** Expression level of each protein in α-pseudoislets on days 0, 5, 10. Data are presented as mean values ± s.e.m. Two-way ANOVA followed by Dunnett’s multiple comparison test. n = 3 biological replicates using islet cells from a single donor **(e,f)**.

We then focused on the genes that were altered over the period of culture exclusively in α-pseudoislets. Among the 263 genes, 110 genes were upregulated and 153 genes were downregulated. We observed that genes including *MT1E* (Metallothionein 1E), *MT1X* (Metallothionein 1X), and *MT2A* (Metallothionein 2A) belonging to the metallothionein gene family and the pathways such as response to metal ions, metallothioneins bind metals, and mineral absorption were upregulated in α-pseudoislets (Fig. 5c,d, Supplementary File 8). Previous studies have identified variant alleles of metallothionein genes and their association with type 2 diabetes (17); however, their role in α-cells is not fully explored. Interestingly, pathways such as FoxO signaling, apoptosis, cell-ECM interactions, cGMP-PKG signaling, MAPK signaling, insulin secretion, and glucagon signaling pathway were all downregulated in the α-pseudoislets (Fig. 5c,d). We observed that the β-cell enriched genes, *DLK1* (Delta Like Non-Canonical Notch Ligand 1) also known as Pref-1, and *GSN* (Gelsolin) were also downregulated significantly over time in α-pseudoislets. On the other hand, *SMIM24* (Small Integral Membrane Protein 24), one of the α-cell enriched genes, was upregulated significantly in α-pseudoislets with time. Using immunostaining, we validated that α-cells express SMIM24, GSN, and DLK1 and expression of these proteins altered with time (Fig. 5e,f). Whether changes in their expression levels are important for α-cells in the absence of cell-to-cell interactions or the paracrine signals derived from other islet cells requires additional studies.

Overall, this study shows that human pancreatic α-cells can be purified by cell dissociation followed by DA-ZP1 labeling and FACS, and the purified α-cells can be maintained in culture to study secretory function, transcriptional changes, survival and proliferation of α-cells.

## Discussion

Despite its importance in both physiology and pathophysiology, the progress in studies related to human α-cell biology has been relatively slower compared to β-cells. A major limitation is the technical challenge in obtaining adequate numbers of high-quality live human α-cells for analyses (18,19). In previous studies, live human α-cells were purified from human islet preparations by using a panel of cell-surface-binding monoclonal antibodies (20) or by combining antibody labeling approaches with ZIGIR labelling (2). Both methods enable isolation of live human α-cells with high purity, but require antibody staining, which increases the risk of cell loss during the labeling procedure and limits α-cell yield. It has been reported that antibody-labeling approaches can recover 200,000 α-cells from 10,000 human IEQ (21). Here we provide an antibody-free approach to obtain primary human α-cells using a fluorescein-tagged sensor which can recover ~ 3x fold higher α-cells with more than 97% purity (Supplementary Figure 1c).

The function of reaggregated α-pseudoislets was examined previously by Liu *et. al*. after sorting individual islet cells by monoclonal surface antibodies (22). In their study, α-pseudoislets secreted glucagon at low concentrations of glucose but failed to respond to changes in glucose concentrations. They argued that the glucose-sensing machinery was deregulated in the absence of paracrine effects of neighboring β-cells. Conversely, our data demonstrated preserved glucose-sensing machinery and secretory actions of α-pseudoislets despite the absence of β-cells (Fig. 3).

While our data suggest that α-cells can regulate glucagon secretion by using cell intrinsic signaling in response to glucose in the absence of paracrine factors, such as insulin and somatostatin (23) and juxtracrine-signaling factors, such as EphA4/7 on α-cells and ephrins on β-cells (15), it cannot exclude their contribution to secretory function *in vivo*. Our RNA-seq data supported maintenance of α-cell identity in α-pseudoislets (Fig. 4a, b), and demonstrated culture time-dependent adaptive alterations in pathways involved in secretory capacity or glucagon signaling (Fig. 5c). Thus, the discrepancy between the two studies could be explained by the use of different methods and time points in performing functional assays.

Human islets are micro-organs composed of multiple endocrine cell types and feature heterotypic cell-cell and cell-matrix interactions. Earlier studies demonstrated that paracrine communication between islet cells are important for their function and proliferation (24). Additionally, vascular endothelial cells, neuronal projections, and ECM network within the islets also contribute regulatory signals to islet cells (25–27). This islet microenvironment is necessarily disrupted when islets are dissociated into single cells as in unsorted islet cells and likely absent in α-pseudoislets. The three groups, namely, i) α-pseudoislets that are bereft of all other endocrine cells, ii) unsorted islets which include all endocrine cells but are distributed in a disorderly manner, and iii) whole islets which consist of all endocrine cells with intact cell-cell interactions allowed us to dissect the effects of the microenvironment on transcriptional regulation of α-cells that reflects paracrine interactions. For example, over the ten days in culture, we observed little change in the transcriptome of whole islets compared with unsorted islet cells (Fig. 4b, Fig. 5a). Although the isolation procedure disrupts innervation and vasculature, the isolated islets preserve cell-cell and cell-matrix interactions. Almost no changes in the transcriptome of whole islets in culture over the period of ten days highlights the importance of intact islet structure on transcriptome stability.

Another example of the significance of the islet cell microenvironment is that genes involved in ECM organization and cell surface interactions were downregulated in unsorted islets cells compared with whole islets (Fig. 4g). These observations are consistent with a previous report showing less abundant ECM components in re-aggregated pseudoislets than in intact islets (28). Accordingly, the mechanosensitive Hippo pathway, essential for integrating cytoskeletal changes with the extracellular environment of the cell was also downregulated in unsorted islet cells reflecting a cellular response to the altered microenvironment (29). Recent studies have revealed Hippo as an important signaling pathway for survival and regeneration of islet cells (30). Indeed, it is known that its downstream protein CCN2 (also known as CTGF) exhibits diverse cell functions including the regulation of β-cell replication in the pancreas during embryogenesis (31).

A notable finding revealed by RNA-seq data is that expression levels of α- or β-enriched genes were recovered after typical islet architecture was regained in unsorted islet cells (Fig. 4b). These data support the fact that intact cell-cell and cell-matrix interactions are important to sustain cell identity *in vitro* (Fig. 4b). The expression levels of α-cell enriched genes did not return towards normal in α-pseudoislets despite their ability to self-assemble and form tight clusters (Supplementary Fig. 2) which might argue that heterotypic cell-to-cell interactions are more important than homotypic interactions in maintaining α-cell identity. Indeed, previous studies reporting an upregulation of β-cell signature genes in α-cells after monotypic α-cell aggregation (32) or extreme β-cell loss (33) support the relevance of heterotypic cell-to-cell interactions in maintenance of α-cell identity. The tighter cluster formation of α-pseudoislets was an interesting observation which could be due to preferential expression of adhesion proteins such as N-CAM (neural-cell adhesion molecule) (34). Overall, our RNA-seq data revealed important transcriptional effects secondary to dissociation of human islets and provide a useful resource for future studies on re-aggregated islet cells.

Among the genes that alter with time in α-pseudoislets, we noted a downregulation of *DLK1*, a positive regulator of insulin synthesis and secretion (35,36). This gene gains relevance considering it is significantly downregulated in type 1 diabetic α-cells (37), and genome-wide association studies identified the *DLK1-MEG3* gene region as a susceptible locus for type 1 diabetes (38). The protein, gelsolin (GSN), which forms a complex with syntaxin 4 protein and induces insulin exocytosis (39) was decreased in α-pseudoislets in a time-dependent manner. This is significant considering gelsolin is among the proteins associated with risk for type 2 diabetes (40). Although we have validated the expression of *DLK1, GSN, SMIM24* that were altered in α-pseudoislets with time; the direct contributions of these proteins to α-cell secretory function in health and disease requires further investigation.

In conclusion, we used a fluorescein-based sensor, to demonstrate isolation of live pancreatic α-cells with high purity and in vitro cultivation of α-pseudoislets without significantly affecting their identity and function over time. This unique tool can be used to isolate live human α-cells for functional studies, transcriptomic analyses, or studies assessing cellular toxicity or proliferation.

## Materials and methods

### Primary Human Cells

Human islets were obtained from the Integrated Islet Distribution Program (IIDP) and Prodo Labs. Upon receipt, islets were centrifuged at 200 g for 1 min and resuspended in fresh Miami medium no. 1A (Cellgro). Cells were transferred to Petri dishes and cultured in 5% CO2 at 37°C. The donor demographic information is summarized in Supplementary File 9. All studies and protocols used were approved by the Joslin Diabetes Center’s Committee on Human Studies (CHS no. 5-05).

### Flow Cytometry

Approximately 15,000 islet equivalents (IEQ) were washed with DPBS and resuspended in TrypLE for single cell dispersion (6). The islets were dispersed into a single cell suspension in TrypLE for 12-15 min at 37°C. Dissociated islet cells were washed with DMEM medium containing 10% FBS and resuspended in the Miami medium containing DA-ZP1 (80 nM) for 10 min at 37°C. The cells were then filtered through a 30 μm filter to remove any aggregates before FACS sorting. DA-ZP1 treated cells were sorted by FACSAria cell sorter (BD Biosciences, Joslin Flow Cytometry Core). Analysis of flow cytometry data was completed using FlowJo 10.7.1 (FlowJo LLC, Ashland, OR). The gating strategy is shown in Figures S1. DA-ZP1 is synthesized by Amit Choudhary’s lab. Compound structure and synthesis of DA-ZP1 are provided in Lee *et. al*. (3) Kahraman *et. al*. (4).

### Immunocytochemistry

Sorted islet cells were spun at 250 g for 5 min, resuspended in Miami media, and immediately seeded in U-bottom low attachment 96 well plates (1,000 cells/200 μl/well or 2,000 cells/200 μl/well). Half of the medium was refreshed every other day. To profile cell types on day 1, the cells were fixed in 4% PFA (Wako) for 15 min at room temperature and washed with PBS. Cells were then permeabilized and blocked with PBS containing 0.25% Triton-X and 5% donkey serum (Sigma) for 1 hour at room temperature. Primary antibodies C-peptide (DSHB, 1:200) and glucagon (MilliporeSigma, G2654, 1:10,000) diluted in antibody dilution buffer (Abcam) were added to the wells for overnight at 4°C. Cells were washed three times with PBS and the secondary antibody, diluted in PBS, was added to the wells for 1 hour at room temperature. Cells were washed three times with PBS and DAPI (Sigma) was added to the wells. Images were captured using an Olympus IX51 Inverted Microscope. For day 5 and day 10, pseudoislets were collected in a tube, washed with PBS, and fixed in 4% PFA for 15 min at RT. The cells were washed, embedded in agarose and paraffin, sectioned and used for immunostaining.

### Cell Viability Assay

The assay was performed using a Cell Titer Glo luminescence cell viability kit according to the manufacturer’s instructions. Approximately eight thousand cells (8 α-pseudoislets, 8 unsorted islet cells, and 8 human IEQ) were added to the wells of opaque-walled 96-well plates and 100 μl of Cell Titer Glo reagent was added to the cells. The content was mixed for 2 min on an orbital shaker and incubated for 10 min at RT. Luminescence was recorded using a Promega GloMax luminometer with an integration time of 0.3 s per well.

### Islet Immunohistochemistry and Quantification

Sections were stained using antibodies against Ki67 (BD550609, 1:100), insulin (Abcam, ab7842, 1:400), glucagon (MilliporeSigma, G2654, 1:10,000 or Abcam, ab92517, 1:5,000) DLK1 (Abcam, ab21682, 1:1,000), GSN (Sigma, HPA054026, 1:50), SMIM24 (Sigma, HPA045046, 1:50), and TUNEL (ApopTag, Chemicon, S7100) and counterstained with DAPI (MilliporeSigma, D9564, 1:6,600). For estimation of islet cell proliferation, 1,000 cell nuclei were counted per section, and data were expressed as percentage of Ki67+ cells. To assess cell death, apoptotic index was measured by quantification of the percentage of TUNEL+GCG+ cells.

### Secretion Assay

Eight α-pseudoislets, 8 unsorted islet cells, and 8 human IEQ obtained from one pancreatic islet donor were handpicked, transferred to wells of a U-bottom 96 well plate, and pre-incubated in Krebs-Ringer bicarbonate (KRB) buffer containing 135 mmol/L NaCl, 3.6 mmol/L KCl, 5 mmol/L NaHCO_3_, 0.5 mmol/L NaH_2_PO_4_, 0.5 mmol/L MgCl_2_, 1.5 mmol/L CaCl_2_, 10 mmol/L HEPES, pH 7.4, 0.1% FFA-BSA) with 16.7 mM glucose for an hour. Static glucose challenge was then initiated by adding KRB buffer containing 3.3 mM or 16.7 mM glucose for 1 h. Aliquots of supernatants were removed for later analysis and ice-cold acid ethanol was added to extract the glucagon content from the cells. Glucagon release and content were measured by the human glucagon ELISA (Mercodia) according to the manufacturer’s instructions.

### RNA isolation, Sequencing, and Data Analysis

Approximately 100 α-pseudoislets, 100 unsorted islet cells, and 100 human IEQ were lysed in TRIzol reagent (Invitrogen) according to the manufacturer’s instructions and the resultant aqueous phase was mixed (1: 1) with 70% RNA-free ethanol. The mixture was added to Qiagen RNeasy micro kit columns and total RNA was extracted following the manufacturer’s protocols. Genomic DNA was digested using RNase-Free DNase kit (Qiagen). The RNA quality and quantity were analyzed using a NanoDrop 1000 Spectrophotometer (Thermo Fisher) and library was constructed using Takara Pico-Input Strand-Specific Total RNA-seq for Illumina (Takara). RNA-seq was performed on an Illumina NovaSeq 6000 according to the manufacturer’s instructions. Approximately 50 million paired-end 100-bp reads were generated for each sample. We aligned the adapter-trimmed reads to the human transcriptome using Kallisto, converted transcript counts to gene counts using tximport, normalized the counts by trimmed mean of M-values (TMM) (7), and transformed normalized counts into log2 counts per million (logCPM) with Voom (8). We applied ComBat-Seq (9) to remove the effect of known batches, and then assessed genes’ association with time and differential expression between groups using the linear regression modeling package limma (10). We corrected for testing many genes with the false discovery rate (FDR). R version 4.1.0 was used. Pathway analysis was done using the ConsensusPathDB interaction database (http://cpdb.molgen.mpg.de/CPDB).

### Singe Cell RNA-seq Analysis of GSE84133

We downloaded this previously published dataset (11) from the Gene Expression Omnibus (GEO). We filtered out cells that have less than 2,000 total gene counts and 1,000 detected genes, and removed genes that have average counts of 0.01 or less. Similar cells were clustered together using a graph-based clustering algorithm and the data was then normalized (12). Moderated t-tests from the linear regression modeling R package limma (10) were performed to detect genes that are differentially expressed between β- and α-cells, with subject and cellular detection rate (i.e. the fraction of detected genes) as covariates.

### Statistical Analysis

All statistics were performed using GraphPad Prism software version 7.0a (GraphPad Software Inc., La Jolla, CA). Specific statistical tests for each experiment are described in the figure legends.

### Study Approval

All human studies and protocols used were approved by the Joslin Diabetes Center Committee on Human Studies (CHS, 5-05).

## Materials Availability

RNA-seq data have been deposited under accession code GSE199412 (The reviewer token is ktwnkokinvopfir). Further information and requests for resources and reagents should be directed to the corresponding author.

## Acknowledgements

We thank Hui Pan and Jonathan Dreyfuss (Joslin Bioinformatics & Biostatistics) for analyzing RNA-seq data, Natalie K. Brown (Joslin) and Oluwaseun Ijaduola (Joslin) for fluorescence microscopy, Alison Marotta (Joslin) and Angela Wood (Joslin) for their assistance with FACS. Flow cytometry experiments were performed in the Joslin Flow Core, supported by the DRC (P30DK036836 and S10 OD021740-01). Human islets obtained from IIDP, supported by NIH (2UC4DK098085), and Prodo Labs.

## Author Contributions

S.K., conceived the study, designed and performed the experiments, analyzed the data, and wrote the manuscript.; K.S. designed and performed cell viability and secretion experiments, analyzed the data, and wrote the manuscript.; D.F.D.J. performed RNA isolation for RNA-seq, J.H. performed immunostaining, D.M. synthesized the DA-ZP1 compound, B.K.W. and A.C. contributed to conceptual discussions, R.N.K. contributed to conceptual discussions, supervised the project, and wrote the manuscript. All the authors have reviewed, commented and edited the manuscript.

## SUPPLEMENTARY FIGURE LEGENDS

**Figure S1.**
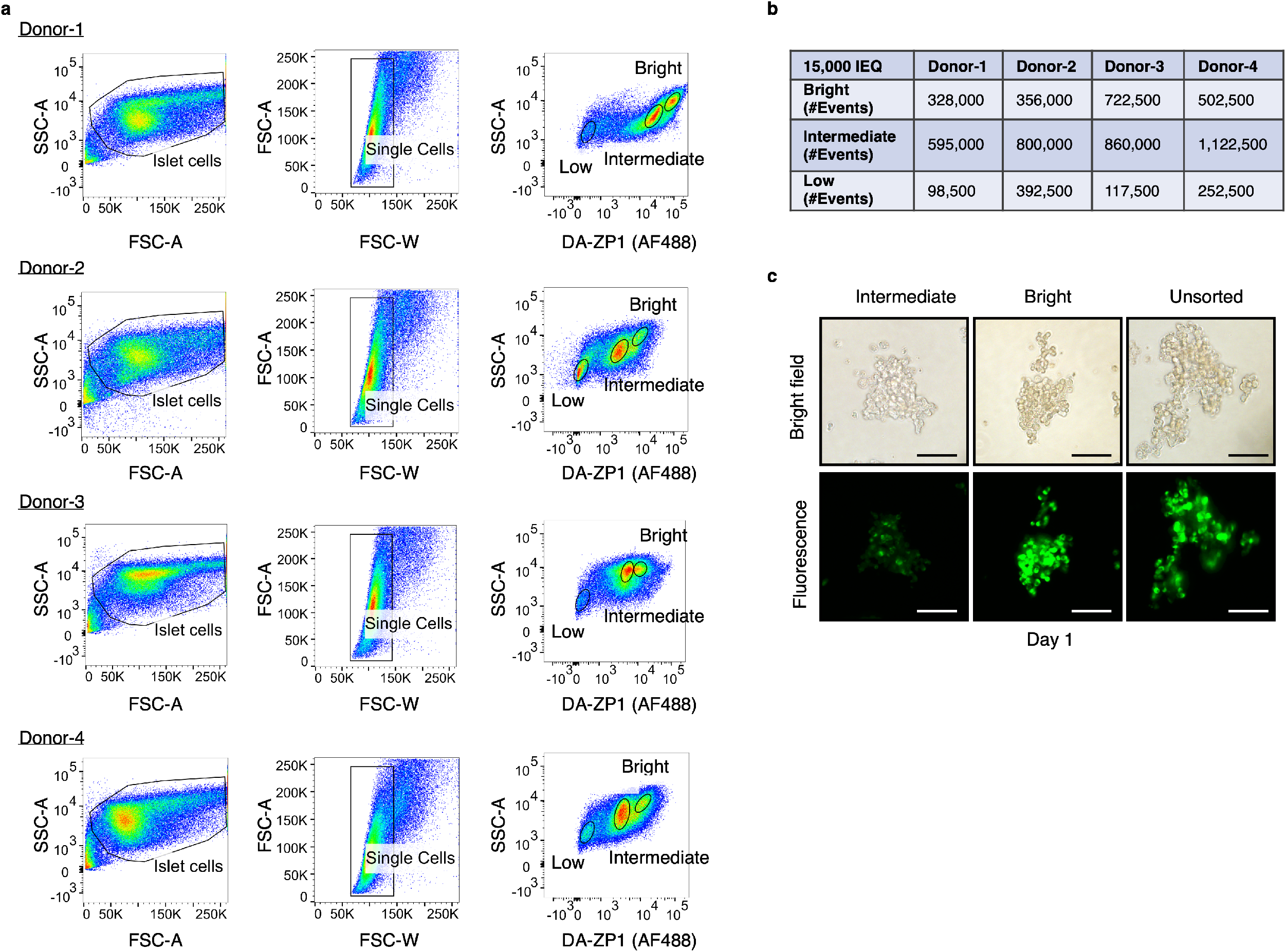
Gating strategy for isolation of live human pancreatic α-cells. **a,** Starting cell population was determined by SSC-A/FSC-A gating. Single human islet cells were gated according to FSC-A/FSC-W gating. Treatment of the single cells with DA-ZP1 resulted in three cell populations with different fluorescence intensity (low, intermediate, and bright). n = 4 human islet donors. Donor information is given in Supplementary File 9. **b,** Number of cells collected by FACS using 15,000 islet equivalent. **c,** The DA-ZP1 derived green fluorescence is maintained in the next day of sorting in the sorted islet cells. The cells were plated in U-bottom well plates to allow re-aggregation after FACS. Scale bar, 100 μm. n = 3 biological replicates using islet cells from a single donor.

**Figure S2.**
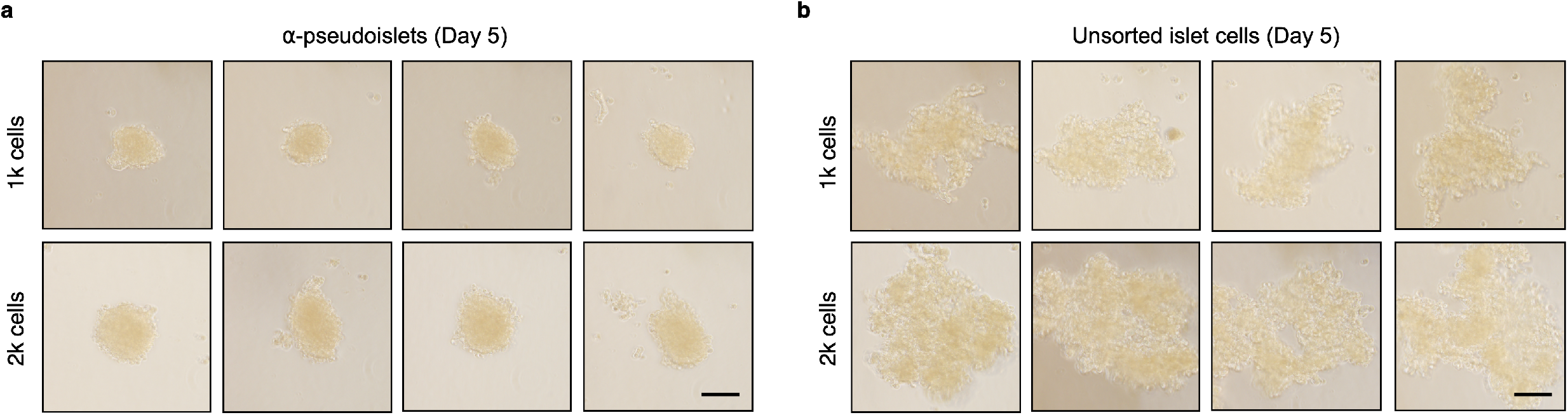
α-Pseudoislets tended to form tighter clusters. Bright field images of sorted α-cells **(a)** and unsorted islet cells **(b)** 5 days post-sorting. 1k (top panel) or 2k (bottom panel) single cells were seeded per well. Scale bar, 100 μm. n = 3 biological replicates using islet cells from a single donor **(a,b)**.

**Figure S3.**
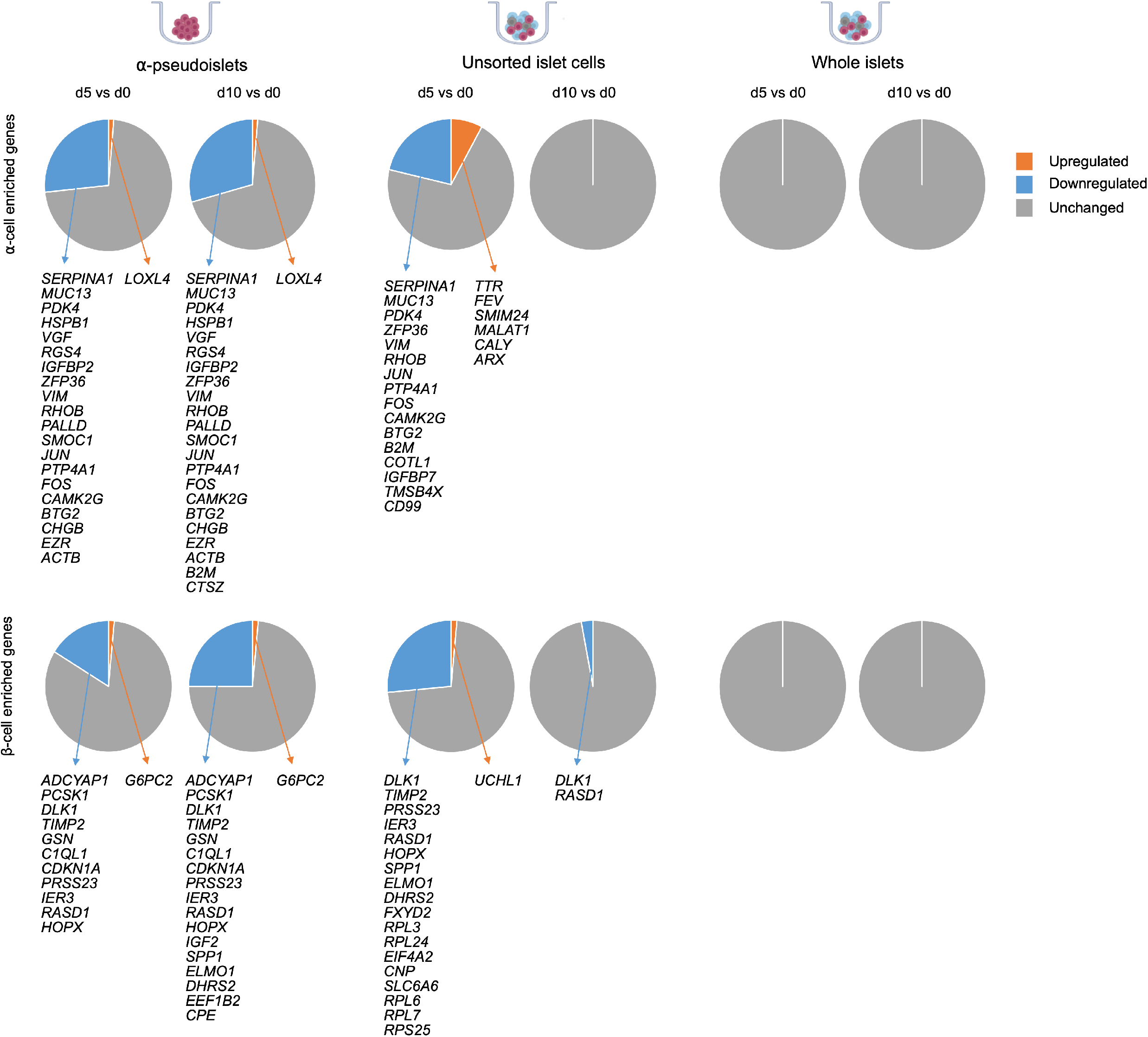
Changes in expression levels of α-cell enriched and β-cell enriched genes in α-pseudoislets, unsorted islet cells, and whole islets on days 0, 5, 10. Pie charts show percentage of genes altered on days 5 or 10 compared to day 0 (FDR < 0.1; FC > 2 upregulated or FC < −2 downregulated). n = 4; α-pseudoislets d0, d5, d10, unsorted islet cells d5, whole islets d5, d10, and n = 3; unsorted islet cells d0, d10, whole islets d0 using islets from four donors.

## References

1. Kirkpatrick CL, Marchetti P, Purrello F, Piro S, Bugliani M, Bosco D, et al. Type 2 diabetes susceptibility gene expression in normal or diabetic sorted human alpha and beta cells: Correlations with age or BMI of islet donors. PLoS ONE. 2010;5(6).

2. Ghazvini Zadeh EH, Huang ZJ, Xia J, Li D, Davidson HW, Li W hong. ZIGIR, a Granule-Specific Zn2+ Indicator, Reveals Human Islet α Cell Heterogeneity. Cell Reports. 2020;32(2):107904.

3. Lee M, Maji B, Manna D, Kahraman S, Elgamal RM, Small J, et al. Native Zinc Catalyzes Selective and Traceless Release of Small Molecules in β-Cells. Journal of the American Chemical Society. 2020;142(14):6477–82.

4. Kahraman S, Manna D, Dirice E, Maji B, Small J, Wagner BK, et al. Harnessing reaction-based probes to preferentially target pancreatic β-cells and β-like cells. Life Science Alliance. 2021;4(4):1–14.

5. Chyan W, Zhang DY, Lippard SJ, Radford RJ. Reaction-based fluorescent sensor for investigating mobile Zn2+ in mitochondria of healthy versus cancerous prostate cells. Proceedings of the National Academy of Sciences. 2014 Jan 7;111(1):143–8.

6. Lee M, Maji B, Manna D, Kahraman S, Elgamal RM, Small J, et al. Native Zinc Catalyzes Selective and Traceless Release of Small Molecules in β-Cells. Journal of the American Chemical Society. 2020 Mar 8;142(14):6477–82.

7. Robinson MD, Oshlack A. A scaling normalization method for differential expression analysis of RNA-seq data. Genome Biology. 2010;11(3):R25.

8. Law CW, Chen Y, Shi W, Smyth GK. Voom: Precision weights unlock linear model analysis tools for RNA-seq read counts. Genome Biology. 2014;15(2):1–17.

9. Zhang Y, Parmigiani G, Johnson WE. ComBat-seq: Batch effect adjustment for RNA-seq count data. NAR Genomics and Bioinformatics. 2020;2(3):1–10.

10. Ritchie ME, Phipson B, Wu D, Hu Y, Law CW, Shi W, et al. Limma powers differential expression analyses for RNA-sequencing and microarray studies. Nucleic Acids Research. 2015;43(7):e47.

11. Baron M, Veres A, Wolock SL, Faust AL, Gaujoux R, Vetere A, et al. A Single-Cell Transcriptomic Map of the Human and Mouse Pancreas Reveals Inter- and Intra-cell Population Structure. Cell Systems. 2016;3(4):346–360.e4.

12. Lun ATL, Bach K, Marioni JC. Pooling across cells to normalize single-cell RNA sequencing data with many zero counts. Genome biology. 2016 Apr 27;17:75.

13. Komatsu H, Cook C, Wang CH, Medrano L, Lin H, Kandeel F, et al. Oxygen environment and islet size are the primary limiting factors of isolated pancreatic islet survival. PLoS ONE. 2017;12(8):1–17.

14. Giuliani M, Moritz W, Bodmer E, Dindo D, Kugelmeier P, Lehmann R, et al. Central necrosis in isolated hypoxic human pancreatic islets: Evidence for postisolation ischemia. Cell Transplantation. 2005;14(1):67–76.

15. Reissaus CA, Piston DW. Reestablishment of glucose inhibition of glucagon secretion in small pseudoislets. Diabetes. 2017;66(4):960–9.

16. Cigliola V, Ghila L, Thorel F, van Gurp L, Baronnier D, Oropeza D, et al. Pancreatic islet-autonomous insulin and smoothened-mediated signalling modulate identity changes of glucagon + α-cells. Nature Cell Biology. 2018;20(11):1267–77.

17. Yang L, Li H, Yu T, Zhao H, Cherian MG, Cai L, et al. Polymorphisms in metallothionein-1 and −2 genes associated with the risk of type 2 diabetes mellitus and its complications. American Journal of Physiology - Endocrinology and Metabolism. 2008;294(5):987–92.

18. Xin Y, Kim J, Okamoto H, Ni M, Wei Y, Adler C, et al. RNA Sequencing of Single Human Islet Cells Reveals Type 2 Diabetes Genes. Cell Metabolism. 2016;24(4):608–15.

19. Brissova M, Haliyur R, Saunders D, Shrestha S, Dai C, Blodgett DM, et al. α Cell Function and Gene Expression Are Compromised in Type 1 Diabetes. Cell Reports. 2018;22(10):2667–76.

20. Dorrell C, Abraham SL, Lanxon-Cookson KM, Canaday PS, Streeter PR, Grompe M. Isolation of major pancreatic cell types and long-term culture-initiating cells using novel human surface markers. Stem Cell Research. 2008;1(3):183–94.

21. Liu W, Kin T, Ho S, Dorrell C, Campbell SR, Luo P, et al. Abnormal regulation of glucagon secretion by human islet alpha cells in the absence of beta cells. EBioMedicine. 2019;50:306–16.

22. Liu W, Kin T, Ho S, Dorrell C, Campbell SR, Luo P, et al. Abnormal regulation of glucagon secretion by human islet alpha cells in the absence of beta cells. EBioMedicine. 2019;50:306–16.

23. Elliott AD, Ustione A, Piston DW. Somatostatin and insulin mediate glucose-inhibited glucagon secretion in the pancreatic α-cell by lowering cAMP. American Journal of Physiology - Endocrinology and Metabolism. 2015;308(2):E130–43.

24. Gromada J, Chabosseau P, Rutter GA. The α-cell in diabetes mellitus. Nature Reviews Endocrinology. 2018;14(12):694–704.

25. Ng XW, Chung YH, Piston DW. Intercellular Communication in the Islet of Langerhans in Health and Disease. Comprehensive Physiology. 2021;11(3):2191–225.

26. Aamodt KI, Powers AC. Signals in the pancreatic islet microenvironment influence β-cell proliferation. Diabetes, Obesity and Metabolism. 2017;19(April):124–36.

27. Lammert E, Thorn P. The Role of the Islet Niche on Beta Cell Structure and Function. Journal of Molecular Biology. 2020;432(5):1407–18.

28. Lorza-Gil E, Gerst F, Oquendo MB, Deschl U, Häring HU, Beilmann M, et al. Glucose, adrenaline and palmitate antagonistically regulate insulin and glucagon secretion in human pseudoislets. Scientific Reports. 2019;9(1).

29. Piccolo S, Dupont S, Cordenonsi M. The biology of YAP/TAZ: Hippo signaling and beyond. Physiological Reviews. 2014;94(4):1287–312.

30. Ardestani A, Maedler K. The Hippo signaling pathway in pancreatic β-cells: Functions and regulations. Endocrine Reviews. 2018;39(1):21–35.

31. Charrier A, Brigstock DR. Regulation of pancreatic function by connective tissue growth factor (CTGF, CCN2). Cytokine & growth factor reviews. 2013 Feb;24(1):59–68.

32. Furuyama K, Chera S, van Gurp L, Oropeza D, Ghila L, Damond N, et al. Diabetes relief in mice by glucose-sensing insulin-secreting human α-cells. Nature. 2019;567(7746):43–8.

33. Thorel F, Népote V, Avril I, Kohno K, Desgraz R, Chera S, et al. Conversion of adult pancreatic α-cells to B-cells after extreme B-cell loss. Nature. 2010;464(7292):1149–54.

34. Cirulli V, Baetens D, Rutishauser U, Halban PA, Orci L, Rouiller DG. Expression of neural cell adhesion molecule (N-CAM) in rat islets and its role in islet cell type segregation. Journal of Cell Science. 1994;107(6):1429–36.

35. Wang Y, Lee K, Moon YS, Ahmadian M, Kim KH, Roder K, et al. Overexpression of Pref-1 in pancreatic islet β-cells in mice causes hyperinsulinemia with increased islet mass and insulin secretion. Biochemical and Biophysical Research Communications. 2015;461(4):630–5.

36. Rhee M, Lee SH, Kim JW, Ham DS, Park HS, Yang HK, et al. Preadipocyte factor 1 induces pancreatic ductal cell differentiation into insulin-producing cells. Scientific Reports. 2016;6(October 2015):1–11.

37. Brissova M, Haliyur R, Saunders D, Shrestha S, Dai C, Blodgett DM, et al. α Cell Function and Gene Expression Are Compromised in Type 1 Diabetes. Cell Reports. 2018;22(10):2667–76.

38. Wallace C, Smyth DJ, Maisuria-Armer M, Walker NM, Todd JA, Clayton DG. The imprinted DLK1-MEG3 gene region on chromosome 14q32.2 alters susceptibility to type 1 diabetes. Nature Genetics. 2010;42(1):68–71.

39. Kalwat MA, Wiseman DA, Luo W, Wang Z, Thurmond DC. Gelsolin associates with the N terminus of syntaxin 4 to regulate insulin granule exocytosis. Molecular Endocrinology. 2012;26(1):128–41.

40. Ngo D, Benson MD, Long JZ, Chen ZZ, Wang R, Nath AK, et al. Proteomic profiling reveals biomarkers and pathways in type 2 diabetes risk. JCI Insight. 2021;6(5).

